# CRB3 and ARP2/3 regulate cell biomechanical properties to set epithelial monolayers for collective movement

**DOI:** 10.1101/2023.04.18.537332

**Authors:** Dominique Massey-Harroche, Vito Conte, Niels Gouirand, Michäel Sebbagh, André Le Bivic, Elsa Bazellières

## Abstract

Several cellular processes during morphogenesis, tissue healing or cancer progression involve epithelial to mesenchymal plasticity that leads to collective motion (plasticity?). Even though a rich variety of EMP programs exist, a major hallmark unifying them is the initial breaking of symmetry that modifies the epithelial phenotype and axis of polarity. During this process, the actin cytoskeleton and cellular junctions are extensively remodelled correlating with the build-up of mechanical forces. As the collective migration proceeds, mechanical forces generated by the actin cytoskeleton align with the direction of migration ensuring an organized and efficient collective cell behaviour, but how forces are regulated during the breaking of symmetry at the onset of EMP remains an unaddressed question. It is known that the polarity complex CRB3/PALS1/PATJ, and in particular, CRB3 regulates the organization of the actin cytoskeleton associated to the apical domain thus pointing at a potential role of CRB3 in controlling mechanical forces. Whether and how CRB3 influences epithelial biomechanics during the epithelial-mesenchymal plasticity remains, however, largely unexplored. Here, we systematically combine mechanical and molecular analyses to show that CRB3 regulates the biomechanical properties of collective epithelial cells during the initial breaking of symmetry of the EMP. CRB3 interacts with ARP2/3 and controls the remodelling of actin throughout the monolayer via the modulation of the Rho-/Rac-GTPase balance. Taken together, our results identified CRB3, a polarity protein, as a regulator of epithelial monolayer mechanics during EMP.

## Introduction

Epithelia are cohesive layers of apico-basally polarized (ABP) cells that adhere and communicate with each other through specialized intercellular junctions and with the substrate through focal adhesions ^1–3^. During development and in some diseases such as cancer, epithelial tissues can perform either complete epithelial to mesenchymal transitions or partial epithelial to mesenchymal transition that has been redefined as epithelial to mesenchymal plasticity ^4–8^. The early phase of EMP mechanically relies on the ability of cells to break their symmetry switching from an apico-basal conformation to a front-rear polarization.

The remodeling of epithelial cells at the periphery of the monolayer requires : i) cell spreading ^9, 10^, by extending basal actin protrusion such as lamellipodia to explore the free space; and, ii) cell migration, by constantly remodeling the focal adhesions that anchor the actin stress fibers at the substrate ^11–17^. During this initiation of migration, a well-balanced coordination of protrusive and retractive forces needs to be maintained by each individual cell that compose the monolayer sheet. Once engaged in the migration process, the leader cells pull via their intercellular adhesions on the follower cells which in turn gradually begin to spread and migrate ^18, 19^. Theoretical and analytical studies suggest that the leader cells might induce normal strain on the follower cells and shear stress on adjacent cells, translating a local stress to coordinated traction forces and cell polarization, which finally results in coordinated motion ^20–22^. So far, many studies have focus on the molecular mechanisms that controls the mechanical forces when the cells are already migrating ^1, 16, 21, 23–28^. Despite extensive studies, the molecular mechanisms that govern the breaking of symmetry of EMP together with the forces underlying this process remains still elusive. In terms of biology during EMP, the remodeling of the cell-cell adhesions, cell matrix adhesions and the reorganization of the actin cytoskeleton correlate with the relocation of the CBR3 polarity complex proteins to the migrating cell front (review in ^12, 29–31)^. The canonical CRB (Crumbs) complex, in mammals is composed of CRB3/Protein Associated with Lin Seven 1 (PALS1) and PALS1-Associated Tight Junction (PATJ) and is known to regulate the actin cytoskeleton organization and intercellular adhesions. CRB is the only protein to possess a transmembrane domain among the polarity complexes, and it has been shown to regulate the organization of intercellular adhesions and actin cytoskeleton in *Drosophila*, zebrafish and mouse embryos ^29^. CRB3 has been shown to play an important function in the establishment and the maintenance of cellular apico-basal polarity ^32–35^, whereas PALS1 and PATJ have been shown to be explicitly involved in the process of collective cell migration ^33, 36–38^. Furthermore, all these data strongly suggest a role for the CRB3 polarity complex in EMP. We still ignore, however, how CRB3 protein and its partners function during the breaking of symmetry of EMP ^39–42^.

Here, we systematically study how CRB3 and its partners impact the epithelial monolayer architecture and mechanics during EMP. We found that CRB3 in tandem with Arp2/3 regulate the breaking of symmetry during the epithelial transition. We have also shown that CRB3 is required for actin remodeling through a permissive effect on Rac1 activation, thereby promoting actin fiber remodeling and alignment along with focal adhesion maturation and orientation and force alignment.

## Results

### Initial breaking of symmetry of EMP leads to actin cytoskeleton remodeling and apical polarity complex relocation

We have developed a new experimental pipeline to reproducibly monitor the biomechanical behavior of an epithelial monolayer transitioning from a static differentiated epithelial morphology to a migratory phenotype. As a cell model system, we adopted the Caco-2 human intestinal cells as they establish a robust ABP when reaching confluency ^43–45^ and actively spread and migrate on free substrates ^46–48^. We used soft lithography to fabricate thin polydimethylsiloxane (PDMS) membranes with a rectangular opening, which we laid on top of a soft polyacrylamide gel substrate coated with collagen-I (Fig. 1A). After 20hr of culture (time t=0 hour), Caco2 cells formed a confluent polarized and symmetric epithelial monolayer and display a cuboidal shape (Fig. 1A, B-E). At this stage, the actin cytoskeleton is organized in well-defined structures running from the apical to the basal domain of the cells. At the apical side of the cells, a dense brush border made by microvilli is observed (Fig. 1 B, red asterisk), along with a lateral cortical actin belt that bridges the cellular junctions (Fig. 1C, green arrowhead). At the basal side of cells, instead, stress fibers are formed (Fig. 1 D) and an actin cable is visible (Fig. 1 D, blue arrowhead) that connects the free cell fronts at the monolayer’s edge. Classically, CRB3, PALS1 and PATJ are mainly located apically at lateral junctions (Fig.1H-J, K,- M, N-0 top left graph, black arrow)^49^.

**Fig. 1:**
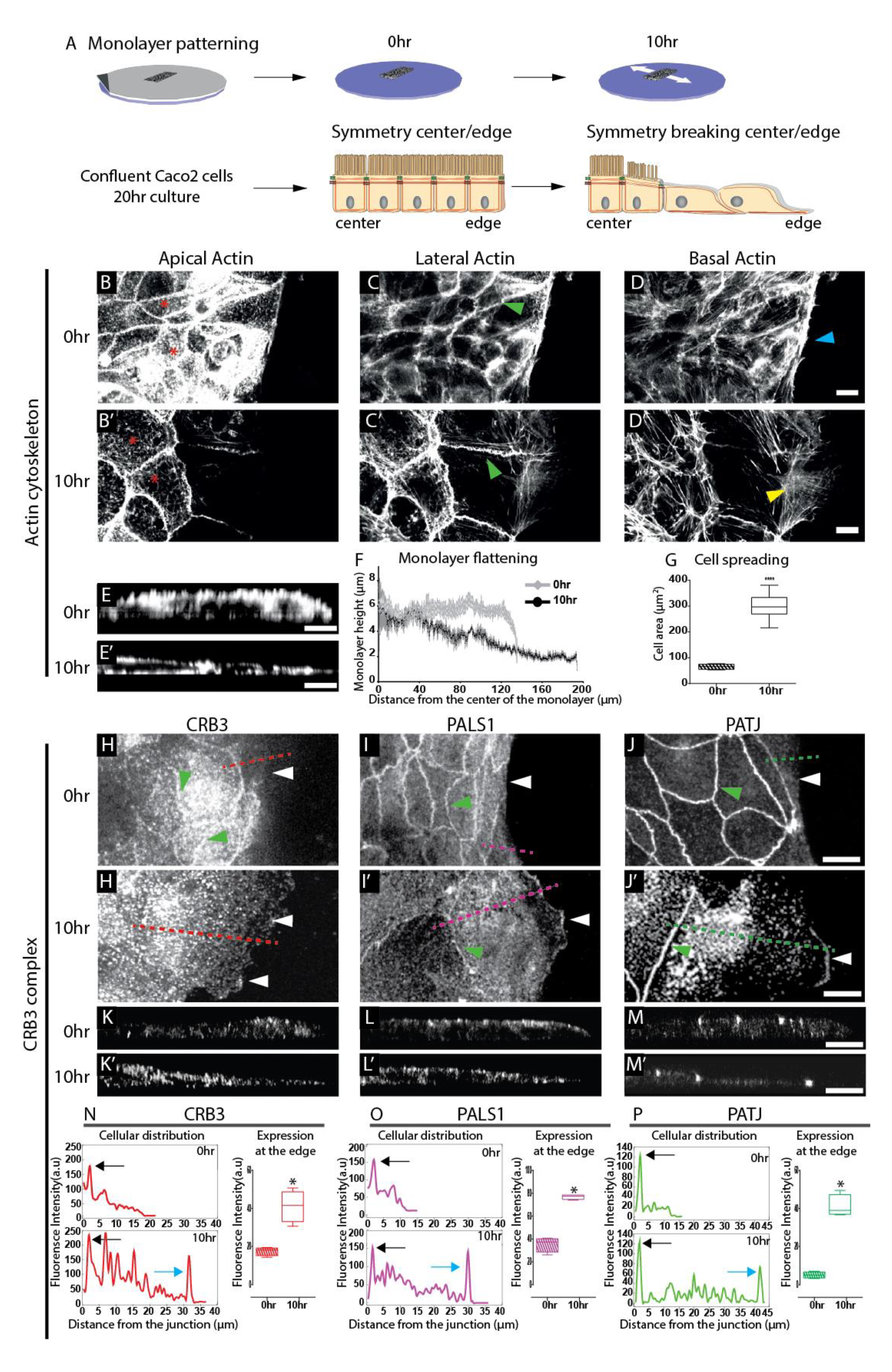
F-actin is remodeled and the polarity complex CRB3 is relocalized during the onset of EMP. A) Scheme of the experimental set up and schematic representation of the organization of the monolayer before and after migration. A PDMS membrane (in grey) is deposited on top of a polyacrylamide gels. Cells are seeded within the rectangular opening and after 20hr the PDMS membrane is removed. Just after removal of the PDMS membrane (0hr) the epithelial cell monolayer is cuboidal and microvilli cover the apical domain, and after 10hr, cells flatten, the microvilli are lost at the edge where lamellipodia form as represented in the drawing. The different domains are represented together with the cellular junction and the actin cytoskeleton. F-Actin organization at 0hr (B-D) and 10hr (B’-D’) of migration. Red asterisks: microvilli, yellow arrowhead: actin arcs, green arrowhead: lateral junctional actin, blue arrowheads: thick actin cable. Lateral view of the F-actin organization at 0hr (E) and 10hr (E’) of migration. Scale bars= 10µm. F) Monolayer height as function of the distance from the center of the cell monolayer. For the different time points several lateral views were measured (0hr, n=44, grey line, 10hr, n= 44, black line). Data are presented as mean ± SEM. G) Quantification of cell area at 0hr (dashed box plot, n=162) and 10hr (empty box plots, n=127) of migration. Data are presented as mean ± Min Max. Localization of CRB3, PALS1, PATJ at 0hr (H-J), and 10hr (H’-J’) of migration. Scale Bar =10µm. White arrowheads point the leading edge of the cell monolayer, green arrowheads point cell-cell adhesions, dashed lines point where the intensity profiles shown in panel N-P are measured. Lateral view of the localization of CRB3, PALS1 and PATJ at 0hr (K-M) and 10hr (K’-M’) of migration. Scale bars= 10µm. N-P) On the left panels, representative fluorescence intensity profile of CRB3 (N), PALS1 (O), PATJ (P) at 0hr and 10hr. Black arrow point the junction of the cell, and blue arrow the peak detected at the edges. On the right panels, quantification of the mean fluorescence intensity of CRB3 (N), PALS1 (O) and PATJ (P) at the edge of the monolayer at 0hr (dashed box plots) and 10hr (empty box plots). For each siRNA 4 fields of views were quantified. Data are presented as mean ± Min Max.

10hr after PDMS stencil removal, the monolayer apico-basal symmetry is broken at free edges, and the epithelial layer flattens (Fig. 1, E’, F) and spreads (Fig. 1 C’, G,). There, cells acquire a migratory state and develop an elongated shape in the epithelial plane. Basally, the actin cable observed at 0hr has vanished, while lamellipodia and actin arcs form (Fig. 1 D’, yellow arrowhead). The data show that the actin cytoskeleton is extensively remodeled during the symmetry breaking, when the free edge cells initiate transition. Moreover, despite the presence of well-defined junctional cortical actin (Fig. 1 C’, green arrowhead), no brush border could be detected, and only sparse microvilli are observed at the apical surface (Fig. 1 B’, red asterisk), thus suggesting a remodeling of the ABP state of free edge cells. Interestingly, actin re-organization correlates with a drastic change in the subcellular localization of the CRB3 polarity complex during EMP. Although PALS1 and PATJ remained located at the lateral junctional belt (Fig1 I’ and J’ green arrowhead, L’, M’, O, P bottom left graph, black arrow), a significant pool became redistributed at the lamellipodial membrane (white arrowhead Fig 1I’, J’ white arrowhead, L’, M’, O, P bottom left graph blue arrow, O, P right graph). In sharp contrast, junctional CRB3 was barely detected at the junction at this stage, and the protein rather accumulated in intracellular vesicles and focal accumulations at the lamellipodia edge (Fig. 1 H’ white arrowheads, K’, N bottom left graph blue arrow, N right graph). These data show that the remodeling of the ABP state of free edge cells triggers the recruitment of the CRB3 polarity complex during monolayer symmetry breaking, albeit with a different distribution between PALS1/PATJ and CRB3. In conclusion, apico-basal symmetry breaking at the free edge of the epithelium triggers the relocalization of the CRB3 polarity complex, which correlates with the remodeling of the actin cytoskeleton,

### CRB3 is required for the epithelial transition to collective migration

In order to test the function of CRB3 polarity complex in epithelial transition, we selectively silenced the expression of CRB3, PALS1 and PATJ using siRNA approach (Fig. S1) and utilized our experimental set up to quantify monolayer migration efficiency (Fig. 2A-E). Phase contrast microscopy revealed that control monolayers (siCT) and siPALS1 knock-down (KD) monolayers are able to migrate efficiently (Fig. 2A, B, E blue and purple curves), with siPALS1 migrating to further extents than the siCT cells (Fig. 2A, B, E). In contrast, depletion of PATJ or CRB3 significantly reduces the monolayer migration (Fig. 2C, D, E green and red curves). Specifically, siCRB3 monolayers exhibit the strongest phenotype with a 2-fold reduction of migration efficiency compared to controls (Fig. 2E, red curve).

**Fig. 2:**
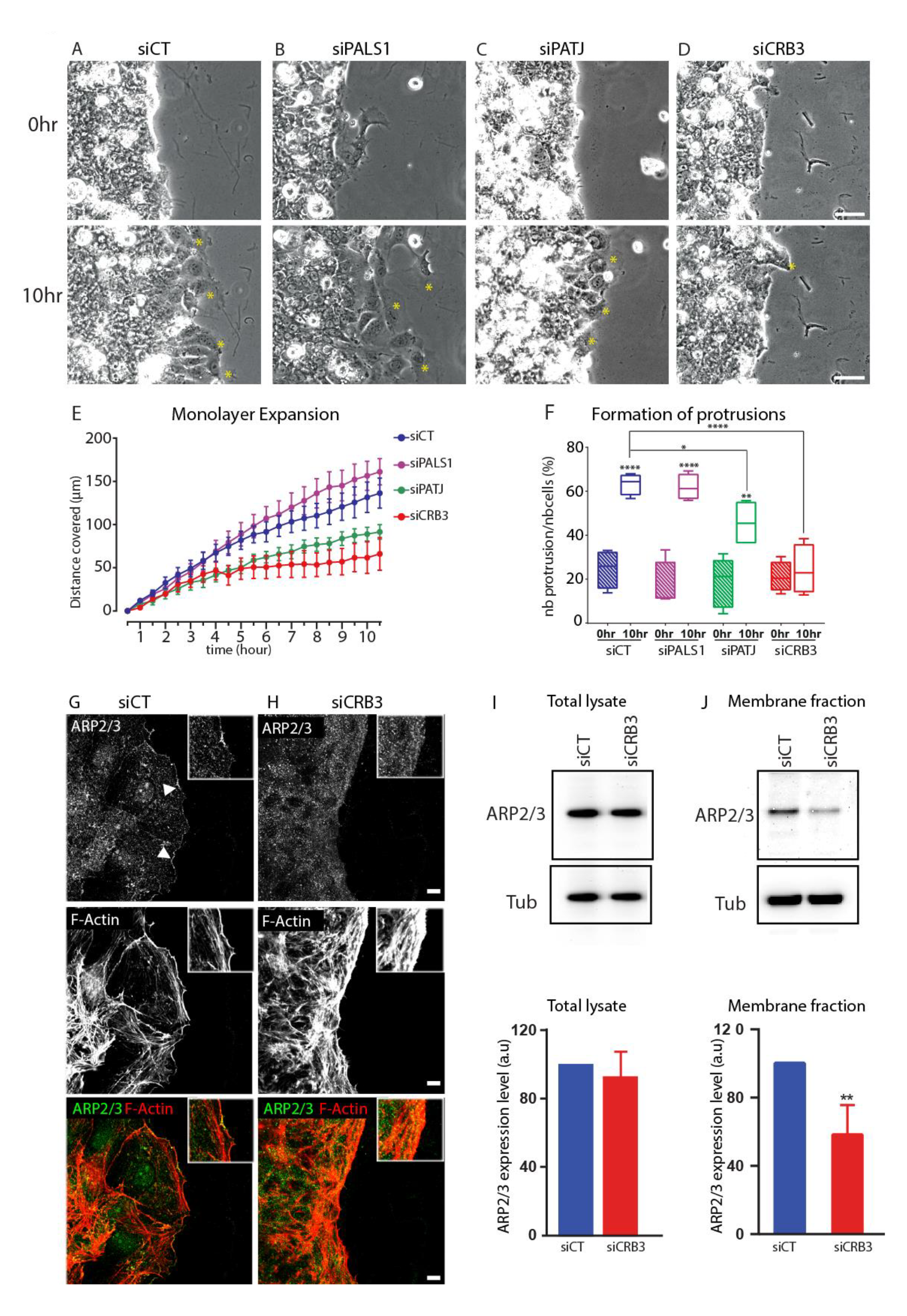
Downregulation of the CRB3 polarity complex alters collective cell migration and the formation of protrusions. Representative phase contrast images showing the effect of each siRNAs (control siCT (A), siPALS1 (B), siPATJ (C) and siCRB3 (D) on cell monolayer expansion at 0hr and 10hr of migration. Yellow asterisks: cell protrusions. Scale bar = 50 µm. E) Time evolution of monolayer expansion (shown as the distance covered by the cell monolayer from the initial point). Number of monolayers for siCT blue, n =7; siPALS1, purple, n =4; siPATJ, green, n=4, siCRB3, red, n=3. F) Cells at the leading-edge bearing protrusions. The box plots represent the ratio between the number of protrusions to the number of cells at the leading edge for each siRNAs at 0hr (dashed box plots) and 10hr (empty box plots) of migration. Number of cells counted for siCT, blue, n=238 0hr, n=228 10hr; siPALS1, purple, n=150 0hr, n=125 10hr; siPATJ, green, n=207 0hr, n=195 10hr; siCRB3, red, n=156 0hr, n=136 10hr. Data are presented as mean ± Min Max. Representative images of ARP2/3 and F-Actin immunostaining in siCT (G), and siCRB3 (H) at 10hr. White arrowheads: edge of the monolayer. Inset are representative zoomed region of ARP2/3 and F-actin immunostainings. Scale bars 10µm. I) Immunoblot analysis showing the expression of ARP2/3 5 days after transfections with siCT and siCRB3. α-Tubulin was used to standardize the loading conditions between the different depletions. Quantification of protein expression levels normalized to siCT cells. Data are represented as mean ± SEM. For each protein, n=3 samples pooled from 3 independent transfections. J) Immunblot analysis of membrane fraction purified lysate of ARP2/3 5 days after transfection with siCT and siCRB3. α-Tubulin was used to standardize the loading conditions between the different depletions. Quantification of protein expression levels normalized to siCT cells. Data are represented as mean ± SEM. For each protein, n=3 samples pooled from 3 independent transfections

In addition, after 10hr, we observed a defect in the cell protrusions at the free edge of depleted monolayers (Fig. 2A-D, yellow asterisks, bottom panels). In the absence of PATJ, a milder phenotype is observed, with 45,7%± 4,2% (Fig. 2F, green undashed box plots) of leading-edge cells forming protrusions, while CRB3 depletion drastically affected the number of membrane protrusions since only 19,1%±3,2% of leading-edge cells bear protrusions (Fig. 2F, red undashed plots). Taken together, these data demonstrate a pivotal role for CRB3 in the breaking of symmetry during epithelial transition, and prompt us to further characterize the cell protrusion defects observed in CRB3-KD monolayers. Analysis of the leading edges at 10hr showed actin filament mis-organization in the absence of CRB3 (Fig.2 G, H, middle panel insets), together with mis-localization of ARP2/3 (Fig. 2 G,H, top panel white arrowheads and inset). In fact, a thick actin cable is formed in CRB3-KD monolayer edges. In addition, ARP2/3 is not further recruited at the protrusion cortex in the mutant free edge cells. Subcellular fractionation and Western blot experiments further confirmed global decrease expression of ARP2/3 associated with the membrane in the absence of CRB3, while the total expression level of ARP2/3 is not affected (Fig.2I, J). Collectively, these results testify of the absence of lamellipodia formation in the absence of CRB3. We thus postulated that the absence of CRB3 may hamper actin remodeling at the cell membrane, preventing the branching of actin and promoting the accumulation of bundled/contractile actin at the monolayer free edge during epithelial transition.

### CRB3 promotes Rac1 activation and actin branching for proper epithelial transition to collective migration

One way to regulate the branched versus the bundled/contractile actin network is by controlling the balance of Rac1 and Rho small GTPases, with Rac1 activation triggering branching of actin via activation of ARP2/3, and Rho activation favoring the bundled/contractile actin ^50, 51^. To measure the level of activation of these small GTPases in CRB3 depleted cells, we performed a GST-pull down assay with GST fused to Rhotekin-p21Binding Domain (GST-Rhot-PBD) or PAK-p21Binding Domain (GST-PAK-PBD) to pull down activated Rho or activated Rac, respectively, in both control and siCRB3 cells (Fig. 3A-C). CRB3 depleted-cells have less activated Rac (Fig. 3 A top panel, B) while Rho activation is increased (Fig. 3 A bottom panel, C), when compared to control cells. These data show a perturbation in the balance of small GTPase activity, with Rho hyperactivation together with Rac hypoactivation in cells depleted of CRB3. This phenotype is compatible with the absence of formation of lamellipodia observed in siCRB3 at the monolayer free edges, and strongly suggest a specific defect in the remodeling of branched actin.

**Fig. 3:**
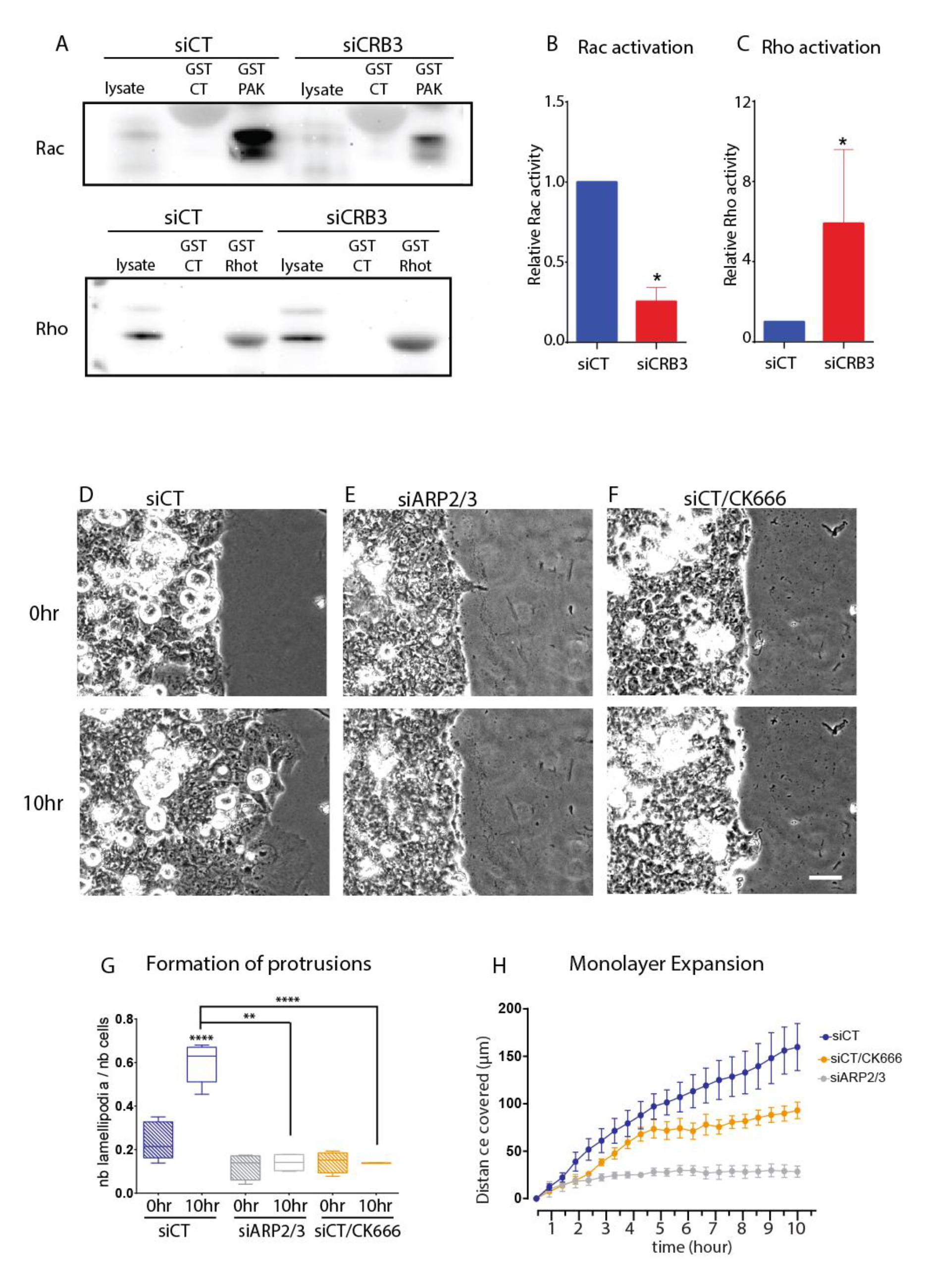
Rho/Rac balance are perturbed in CRB3 depleted cells, and inhibition of branched actin mimic siCRB3 phenotype. A) Representative western blot images showing the effect of CRB3 downregulation on the activation of Rac (top panel) and Rho (bottom panel). siCT and siCRB3 lysates were pulled-down with GST-CT, GST-PAK-PBD (top panel) and GST-Rhot-PBD (bottom panel) and probed to detect Rac (top panel) and Rho (bottom panel) expression. The activation of Rac (B) and Rho (C) were quantified by computing the ratio between the amount of Rac and Rho pull-down to the total expression detected in the siCT and siCRB3 lysates. The data presented in the histograms are the mean ± SEM of 3 independent experiments for siCT and 3 independent experiments for siCRB3. D-F) Representative phase contrast images showing the expansion of siCT cells, siARP2/3 and siCT/CK666 treated cells at 10hr of migration. Scale bar = 50 µm. G) Cells at the leading-edge bearing protrusions. The box plots represent the ratio between the number of protrusions to the number of cells at the leading edge for each siRNAs at 0hr (dashed box plots) and 10hr (empty box plots) of migration. Number of cells counted for siCT, blue, n = 123 0hr, n =126 10hr; for siCT/CK666 treated cells, orange, n =165 0hr, n =152 10hr; for siARP2/3 n=114 at 0hr, n =120 at 10hr. H) Time evolution of monolayer expansion (shown as the distance covered by the cell monolayer from the initial point). Number of monolayers for siCT blue, n =3; siCT/CK666, orange, n =3; siARP2/3, gray, n=3.

We further assessed whether a direct link exists between branched actin remodeling and symmetry breaking during epithelial transition. As expected from literature^52, 53^, inhibiting actin branching with CK666 drug treatment or using siARP2/3 transfection blocks the formation of cell protrusion and prevents the global migration of the epithelial control monolayer (Fig. 3D-H). These results nicely mimicked the CRB3 depletion phenotype we observed in Figure 2. In addition, ARP2/3 has been shown to directly interact with CRB3 in Sertoli cells^54^. We thus investigated whether CRB3 binding to ARP2/3 also takes place in our system. By using affinity-precipitation (peptide pull-down) with the full-length cytoplasmic domain of CRB3 (90-120 AA, CRB3 cyt) or the same cytoplasmic domain only containing the FERM binding domain (90-100 AA, CRB3FERMBD) that binds to cytoskeletal associated proteins such as ARP2/3 ^54^ (Fig. S2A), we revealed that ARP2/3 interacts with CRB3-FERM binding domain construct (Fig. S2B).

To determine whether a functional correlation exists between CRB3 and ARP2/3 for branched actin remodeling, we scrutinized actin fiber arrangement in siCT, siCRB3 and siARP2/3 cells at 0hr and 10hr after removing the PDMS stencil (Fig 4). Whereas CRB3 or ARP2/3 silencing did not affect the initial organization of actin cytoskeleton (Fig. 4 A-C, S3), differences appear between siARP2/3, siCRB3 cells and siCT cells after 10hr of PDMS removal (Fig. 4 A’-C’, S3). In a similar manner, both siCRB3 and siARP2/3 cells do not exhibit lamellipodia but instead still display a thick actin cable at their free basal edge (Fig 4 B’,C’ blue arrowhead). These data clearly state that the phenotypes observed result from a defect in the disassembly and remodeling of the actin cytoskeleton. Further segmentation of the actin filaments and quantitative analysis of their organization in the basal cell leading edge revealed that no clear difference could be noted between the 3 conditions at 0hr (Fig 4D-F, G-H dashed boxes). At 10hr, however siARP2/3 cells developed a thick actin cable oriented orthogonally to the migration direction (Fig 4 F’, blue inset), as siCRB3 cells did (Fig.4E’, blue inset). In addition, quantification of alignment coherence and orientation index, where 0 reflects an orientation of the fibers in the direction of migration and 1 reflects an orientation of the fibers perpendicular to the direction of migration (see Materials and Methods), shows a reduced orientation index and less coherent alignment of actin fibers in siCRB3 and siARP2/3 at 10h compared to control cells (Fig.4 E’, F’, G, H undashed boxes). These quantifications reflect a loss of the polarized state of the monolayer at the free edge in siCRB3 and siARP2/3 cells. All together, these data show that CRB3 and ARP2/3 are both needed for the correct organization of the actin fibers tangentially to the direction of migration. As remodeling of the actin cytoskeleton is coupled to changes in cell shape, it can be thus considered as the deformability potential of cells ^55, 56^. We therefore computed the cell area, height and use the changes in cell area as a proxy to compute the strain rate of cell spreading (Fig. S3). Control cells presented a high rate of deformation in spreading with a strain rate of 4,35 demonstrating that Caco-2 cells strongly remodel their shape when they start to migrate at 10hr (Fig S3K). In sharp contrast, a quasi-null strain rate of 0,89 and 1,16 was computed for siCRB3 and siARP2/3 cells respectively (Fig S3K), siCRB3 and siARP2/3 cells being taller and expanding less than controls (Fig. S3A’-I’, J undashed box, M). Our data shows that CRB3 together with ARP2/3 are essential for the remodeling of epithelial cell shape, and in particular for cell spreading, during the initial transition from epithelial to migrating cells through the remodeling of the actin cytoskeleton.

**Fig. 4:**
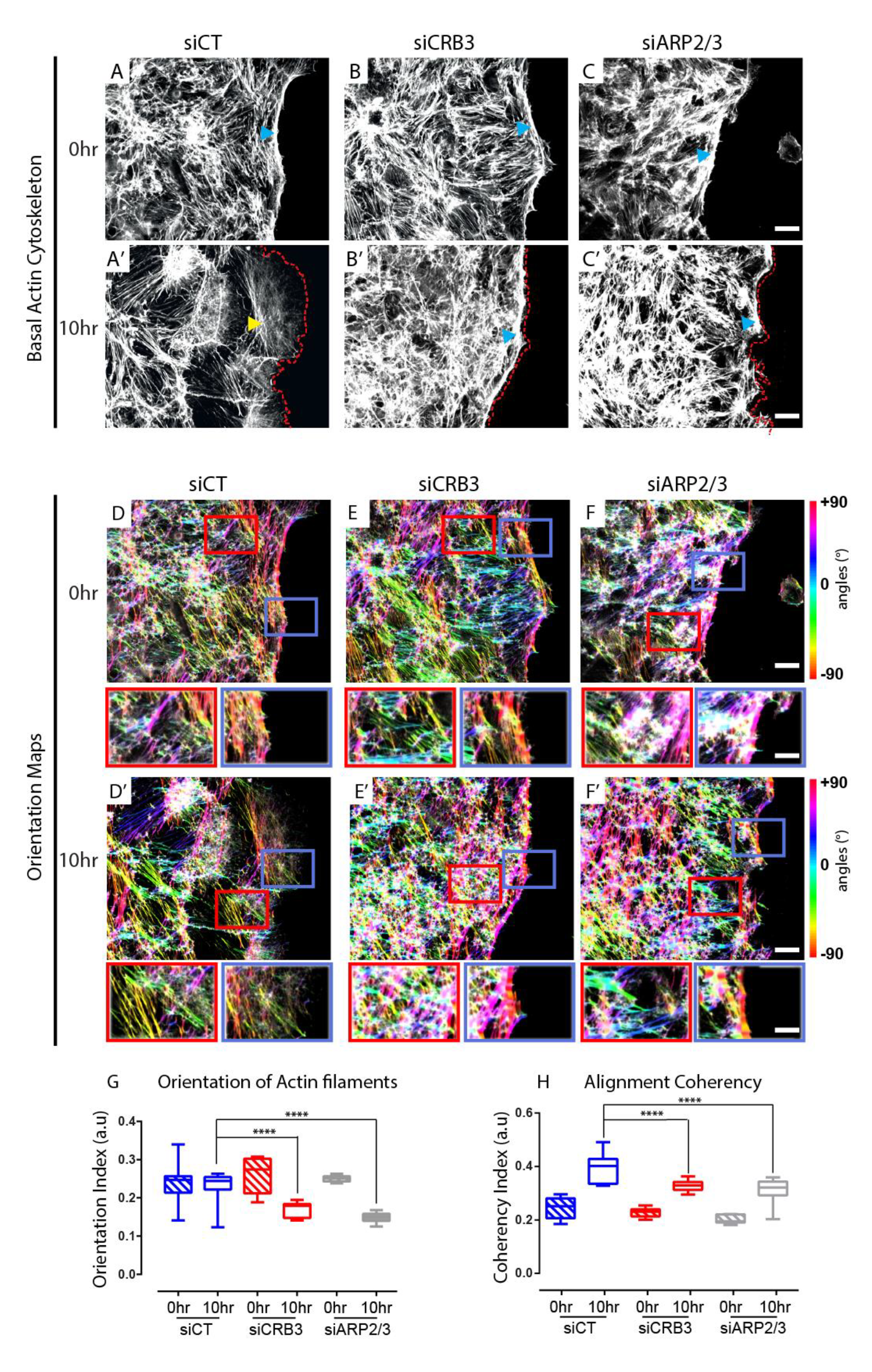
Downregulation of the CRB3 complex and ARP2/3 alter the organization of the F- actin. A-C’) Representative images showing the effect of each siRNA on the organization of the F-actin at the basal domain (siCT (A, A’), siCRB3 (B, B’), siARP2/3 (C, C’ (at 0hr and 10hr of migration. Blue arrowheads: F-actin cable, yellow arrowheads: actin arcs, red dash line: outline for the basal edge. Scale bars= 15 µm. D-F’) Representation Orientation maps of actin fibers (siCT (D,D’), siCRB3 (E,E’) and siARP2/3 (F,F’). Scale bar in degree. Insets are representative zoomed regions of actin orientation. G) Quantification of actin fibers orientation area for each siRNAs at 0hr (dashed box plots) and 10hr (empty box plots) of migration. For each siRNA 10 fields of views were quantified. siCT, blue; siCRB3, red; siARP2/3, grey. Data are presented as mean ± Min Max. H) Quantification of actin fibers coherency for each siRNAs at 0hr (dashed box plots) and 10hr (empty box plots) of migration. For each siRNA 10 fields of views were quantified. siCT, blue; siCRB3, red; siARP2/3, grey. Data are presented as mean ± Min Max.

In summary, using a variety of tools, we have shown that CRB3 is required for cells to properly activate Rac1 and remodel the actin cytoskeleton in a similar way to the actin branching protein ARP2/3.

### CRB3 and ARP2/3 regulate the remodeling of cell-matrix adhesions

The remodeling of the actin cytoskeleton is required for the cell-substrate adhesion (review in ^57^). We therefore decided to study the focal contacts in our system, to decipher a potential correlation between the organization of the actin filaments and the organization of the focal adhesions. Focal adhesions were immunostained for paxillin, at 0hr and 10hr after removal of the PDMS membrane in the different depletions (Fig. 5A-C, A’-C’), and image segmentation and analyses of the focal contacts were performed to measure their maximal length, size repartition orientation (Fig. S4, Fig.5 G, H) and spatial dispersion of focal contacts (Fig.5D-F, D’-F’, I). Moreover, as previous works have shown that, as cells start to migrate, their focal contacts are remodeled and maturated from small nascent adhesion to larger focal adhesions ^58, 59^, we subdivided accordingly the focal adhesions into three different categories: nascent <0.25µm, focal complex <0.5µm, focal adhesions >1-5µm ^57^. Therefore, by looking at the ratio of the different adhesion categories at 0hr and 10hr, we addressed whether the focal contacts are able to be remodeled and matured from nascent to focal adhesions. At 0hr, no significant difference was measured in the different conditions, indicating that focal adhesions were not affected by any of the depletions when the cells are in a static epithelial state (Fig. 5A-C, G-H, dashed boxplots). At 10hr, an increase in size with a higher ratio of mature focal adhesions was observed in siCT cells (Fig 5G, S4G). As siCT cells spread, the length of the actin filament increased and this was translated by an increase in the dispersion of the focal adhesion (Fig 5 D-D’, I undashed blue box). At the opposite, siCRB3 and siARP2/3 cells exhibited an opposite trend with smaller immature focal contacts (Fig. 5G, undashed boxplot), and a significant decrease of their orientation and dispersion (Fig. 5 H-I, undashed boxplot) in comparison with control cells.

**Fig. 5:**
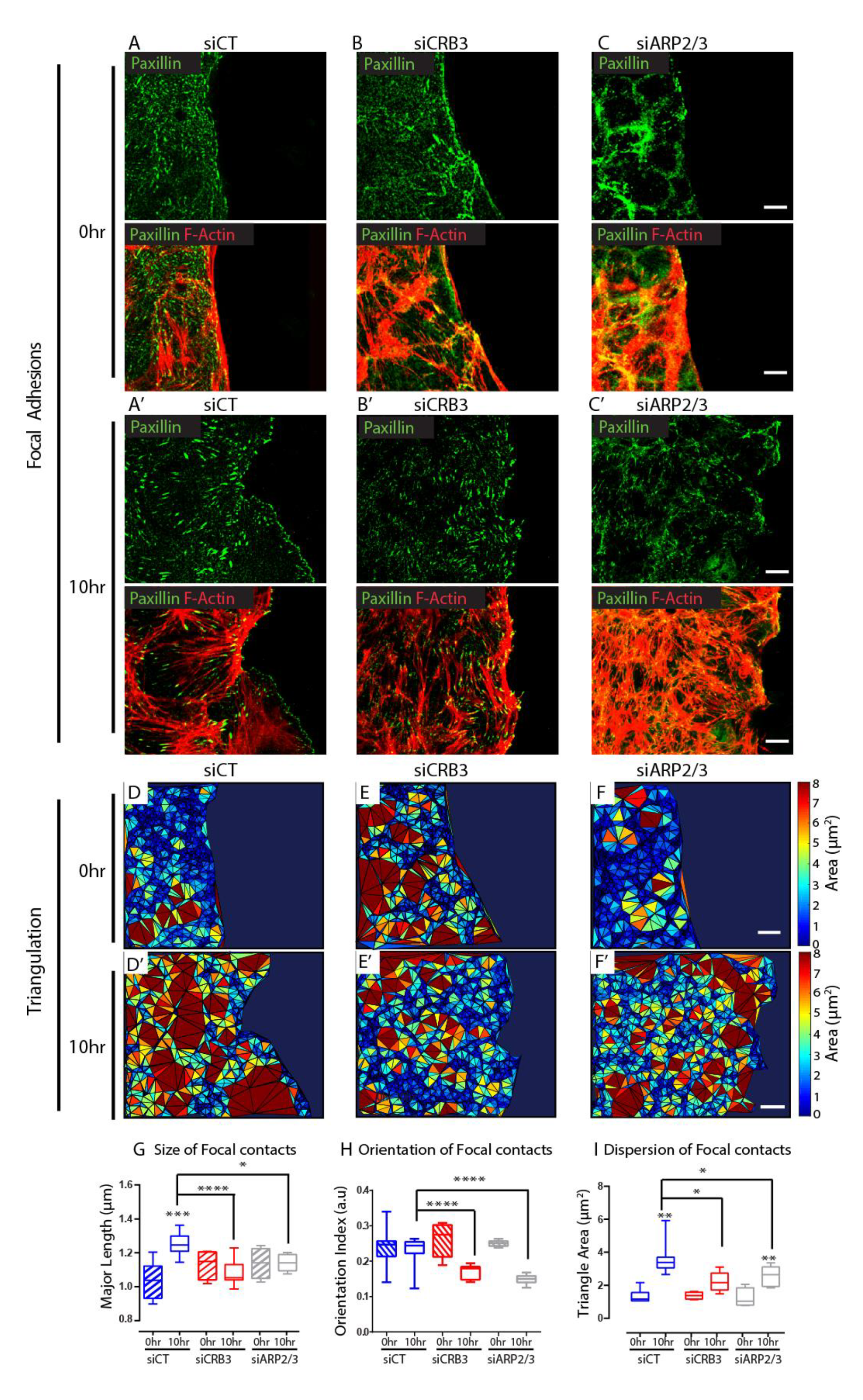
Downregulation of the CRB3 and ARP2/3 alters the organization of the focal contacts. Representative images of focal adhesions immunostained with paxillin (top) and paxillin/F-actin (bottom) for siCT(A), siCRB3 (B), siARP2/3 (C), at 0hr and siCT (A’), siCRB3 (B’), Arp2/3 (C’) at 10r.. Scale bars 10µm. Representative maps of the Delaunay Triangulation of paxillin for siCT (D), siCRB3 (E), siARP2/3 (F) at 0hr and siCT (D’), siCRB3 (E’), siARP2/3 (F’) at 10hr. Scale Bar 10µm. G) Quantification of the major length of the focal contacts is represented as dashed box plots (0hr) and empty box plot (10hr). Data are represented as mean ± Min Max. H) Quantification of the orientation of the focal contacts is represented as dashed box plots (0hr) and empty box plot (10hr). Data are represented as mean ± Min Max. The total number of focal contacts quantified for siCT, blue, n = 2530, 6 fields of view, 0hr, n= 6652, 15 fields of view, 10hr; siCRB3, red, n=2230, 4 fields of view, 0hr,n=7410, 17 fields, 10hr; siARP2/3, grey, n=1476, 4 fields of view, 0hr, n =5806, 13 fields of view, 10hr. Data are presented as mean ± Min Max. I) Quantification of the area of the triangle obtained by a Delaunay triangulation over all the focal contacts for each siRNA condition at 0hr (dashed boxplot) and at 10hr (empty boxplot). The box plots represent the mean ± Min Max of all the triangles measured for each siRNAs. siCT, blue, n=23232, 7 fields of view, 0hr, n= 25140, 16 fields of view, 10hr; siCRB3, red, n= 12917, 4 fields of view, 0hr, n=37164, 18 fields of view, 10hr, siARP2/3, grey, n= 11737, n=4, 0hr, n= 13816, 6 fields of view, 10hr.

In conclusion, defects in shape, orientation and distribution of focal adhesions in siCRB3 and siARP2/3 cells showed the requirement of CRB3 together with ARP2/3 in remodeling focal contacts, and may be explained by a defective remodeling and growth of the basal actin fibers.

### Loss of CRB3 or ARP2/3 perturbs the mechanics of the cellular monolayer

Actin dynamics and focal adhesions are related to force generation and transmission across epithelial cell monolayers ^31, 60^. We thus postulate that CRB3 together with ARP2/3 could be involved in the generation and alignment of forces that are required for the cells to initiate efficient collective motion ^20–22^.

Traction force microscopy and monolayer stress microscopy ^20, 21, 23^ were used to measure forces that cells exert at the surface of the substrate. To visualize the orientation and alignment of tension, we plot the vectorial fields of tension. At 0hr, before the symmetry breaking, tractions exhibited a punctate pattern with higher tractions forces being exerted toward the edge of the monolayer in control cells as in siCRB3 and siARP2/3 cells, with depleted cells exhibiting overall higher tractions (Fig.6D-F). In a similar manner, tensions are higher in siCRB3 and siARP2/3 cells compared to control cells (Fig. 6 G-I) and exhibit lower anisotropy as represented by the orientation of maximal force tensor (Fig. 6J-L). After 10hr, traction forces and tension increase with cells depleted for CRB3 and ARP2/3 exerting higher forces in comparison with controls (Fig.6D’-F’, G’-I’). Interestingly, the anisotropy of the maximal force tensor increases as the vectors tend to align toward the direction of migration in control cells (Fig 6J’). In contrast, in siCRB3 and siARP2/3, we still observed strong anisotropy and disorganization of the vectors at 10hr (Fig. 6K’, L’). To give a comprehensive understanding on how the mechanics evolved in response to the different siRNAs, we computed the temporal evolution of tension, tractions, and of the Shannon’s entropy in the vectorial fields. This analysis revealed that control cells, siCRB3 and siARP2/3 cells increased their tractions and tension, albeit forces in siCRB3 and ARP2/3 being always higher than in control cells (Fig.6M, N). Interestingly, cells depleted for CRB3 and ARP2/3, are not able to align their tension as the control cells do, as shown by the increased entropy in siCRB3 and siARP2/3 cells (Fig.6 O). Thus, depletion of CRB3 or ARP2/3 impacted the global mechanical properties, intensity and organization, of the cell monolayer.

**Fig. 6:**
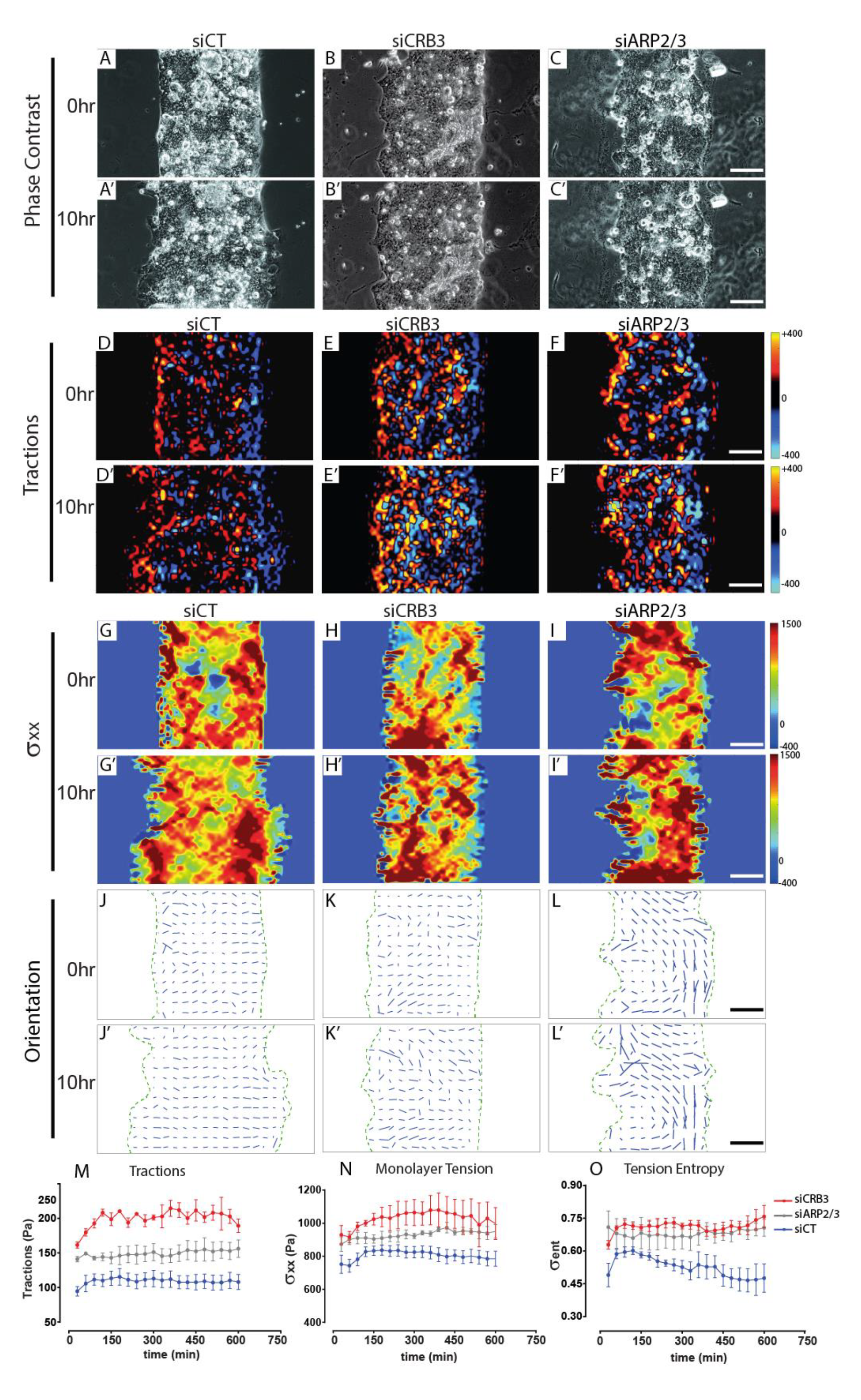
Downregulation of the CRB3 complex and Arp2/3 alters monolayers physical properties. Representative maps showing the effect of each siRNAs on monolayers dynamics at 0hr and 10hr of migration. Representative phase contrast images of siCT (A), siCRB3 (B), siArp2/3 (C) at 0hr and siCT (A’), siCR3 (B’), siArp2/3 (C’) at 10hr. Representative traction force maps for siCT (D), siCRB3 (E), siARP2/3 (F) at 0hr and siCT (D’), siCRB3 (E’), siARP2/3 (F’) at 10hr. Representative intercellular tension maps σ_xx,_, or siCT (G), siCRB3 (H), siARP2/3 (I) at 0hr and siCT (G’), siCRB3 (H’), siARP2/3 (I’) at 10hr. Representative vectorial fields of intercellular tension for siCT (J), siCRB3 (K), siARP2/3 (L) at 0hr and siCT (J’), siCRB3 (K’), siARP2/3 (L’) at 10hr.. Scale bars= 100 µm. M-O) Time evolution of traction forces (M)intercellular tension (N), tension entropy (O) for the siCT (blue), siCRB3 (red), siARP2/3 (grey). Data are presented as mean ± SEM. siCT n=7 independent cell monolayers, siCRB3 n=3 independent cell monolayers, siARP2/3, n = 3 independent monolayers.

To further determine whether a correlation occurs between biological and mechanical phenotypes, we selected morphometric parameters (cell area, cell orientation, cell elongation, focal adhesion area, focal adhesion orientation, focal adhesion elongation, actin fiber orientation and coherency) and mechanical parameters (monolayer kinetics, tractions, tension and entropy) and summarized our data in two matrices (Fig. 7), 0hr and 10hr, for three conditions, (Fig. 7A-D). We computed two matrices in which each element contains the z-score of the morphometric and mechanical properties in response to each depletion. This numerical analysis demonstrates that the mechanical parameters are markedly affected at 0hr for both KD cell lines, while a relevant impact is not measured in the morphometric parameters. At 10hr, we observed a reinforcement of the mechanical properties along the same trends observed at 0hr (Fig. 7B, D), while drastic modifications can be observed for the morphometric properties (Fig. 7A, C).

**Fig. 7:**
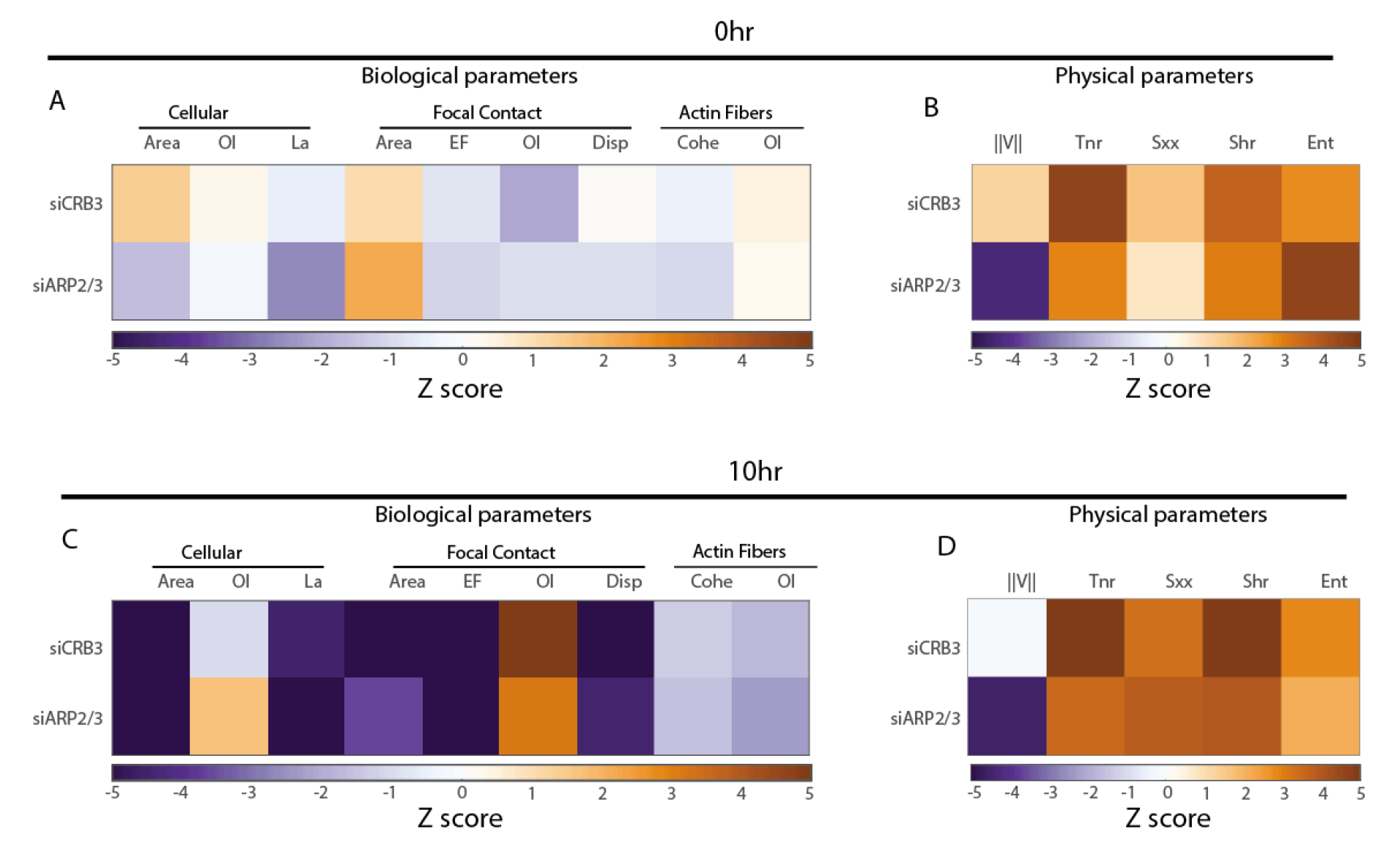
The depleted cells exhibit different mechanical properties before showing difference at the biological level. Effect of siRNAs on biological and physical properties expressed in z-scores at 0hr (A, B) and at 10hr of migration (C, D). OI: Orientation Index, La : number of lamellipodia, EF : Elongation Factor; Disp :Dispersion, Cohe : Coherency, V : velocity, Tnr : normal Traction, Sxx : Normal component of the stress tensor in the x direction, Shr : Shear component of the stress tensor, Ent: Entropy.

Collectively, these data demonstrated the reciprocity between cell behavior and mechanical features, and in particular, the relationship between the remodeling of actin cytoskeleton/focal adhesions, and the loaded forces the cells exert at the cell-cell interface and on their underlying substrate.

## Discussion

Epithelia transition towards collective migration is associated with changes in cell polarity and adhesion in epithelial tissues. The transition from a static monolayer to collective migration begins with the breaking of symmetry from apical-basal to front-rear at the monolayer edge and ends up with collective cell motion. These steps require a coordinated remodeling of intercellular adhesions, focal contacts and actin cytoskeleton. In our study we have implemented a system that allows the systematic study the transition from a static monolayer where epithelial cells polarize in the ABP axis, to a more dynamic monolayer where leading edge migrating cells develop instead front-to-rear polarized state.

Here we have identified CRB3 / ARP2/3 as mechanoregulatory module of the symmetry breaking during EMP. Several studies performed on different cellular models, such as non-tumorigenic human mammary epithelial cells, tumorigenic kidney-derived cells or colorectal adenocarcinoma cells have led to conflicting conclusions regarding the function of CRB3 during collective cell motion ^39–42^. In human mammary epithelial cells or tumorigenic kidney cells, the loss of CRB3 expression increases cell invasion, and promotes cell scattering while our study together with the work of Lioka et al^39^ clearly demonstrates that loss of CRB3 prevents cell spreading. The discrepancies between different cell lines might be explained by different factors. Firstly, the regulation of protein expression such as CRB3 depends on the model system used ^40, 61–64^. Recently, it has been described that CRB3 gene expression differs depending on the tumoral tissues or cell models ^39, 61–63^. As an example, CRB3 is upregulated in colorectal and breast cancer whereas in glioblastoma it is downregulated. In the study of Mao et al^63^, the authors also described that in non-tumorigenic breast MCF10A cells, CRB3 expression and localization depends on the cellular density with a lower expression in confluent cells when compare to sparse cells. These different expression patterns could result in the formation of distinct complexes, and trigger various signaling pathways that could lead to a great variety of phenotypes. These differences should be considered before raising a conclusion regarding the role of CRB3 during collective cell migration. Secondly, the choice of the cellular model is crucial to address properly the question of the initial step of the transition from a static differentiated epithelial to a migratory epithelial monolayer, when cells dramatically change their polarity and shapes as well as their actin organization. Caco2 cells are highly polarized with a dense apical actin network, forming a brush border, and robust cellular adhesions, mimicking i*n vivo* enterocyte organization. When cells start to spread, the polarity proteins are relocalized, the apical actin network is disassembled and cellular adhesions are remodeled, leading to a dramatic cell shape change. The study by Li et al, used MCF10A mammary epithelial cells, which are not columnar, do not exhibit a dense apical actin brush border and have weaker cellular adhesions ^61, 64, 65^. In the case of MCF10A cells, the remodeling of the actin cytoskeleton is less drastic when compared to Caco2 cells, as the cells do not undergo a massive change of shape. Therefore, the results obtained with this MCF10A cell line cannot be generally transposed to understand the breaking of symmetry during EMP. The Caco2 cell line used in the present study is a more suitable epithelial model to address the regulation of the actin cytoskeleton during the initial cell spreading and allows us to unveil a key role of CRB3 in this process.

Our study was, however, not limited to CRB3 but also identified new functional module for EMP composed by CRB3 and ARP2/3. By performing a global quantitative analysis of several biological and mechanical parameters, we were able to demonstrate with an unbiased approach that CRB3 and ARP2/3 exhibit similar functions and could be part of the same functional module during EMP. In line with this conclusion, our biochemical data also show that CRB3 can bind to ARP2/3 independently of PALS1, confirming a heterogeneity of CRB3 complex composition. Using Caco2 cells, we describe ARP2/3 as a potential mechanobiological effector of CRB3, and show that CRB3 expression regulates the small GTPase Rac/Rho balance. CRB3 is essential for the cells to activate Rac1, and with ARP2/3 they promote the remodeling of actin and the maturation focal adhesion during the breaking of symmetry of EMP. Previous studies have shown a role of the CRB polarity protein complex in the regulation of the actin cytoskeleton in other cellular processes ^29^ and here we have found a new cellular context for the role of CRB3 in regulating actin organization. CRB3 is a small transmembrane protein that forms a polarity complex with PALS1 and PATJ which can both recruit regulatory proteins to modulate actin organization. In our study we show, however, that CRB3 can interact with ARP2/3 independently of PALS1 and PATJ, via its FERM binding domain. Our data suggests that to break epithelial symmetry, CRB3 might be relocated to the leading edge where, as a transmembrane protein, it recruits ARP2/3, an actin binding protein that is necessary for the formation of protrusions. This localization leads to cell polarization with a polarized activation of the Rho/Rac balance in a rear-to-front fashion depending on the localization of CRB3 as proposed by several studies ^18, 29, 66^. During this initial cell spreading, this local activation of Rac induces cytoskeletal rearrangements with a rapid actin polymerization and alignment of the actin filament promoting the engagement and maturation of focal contacts ^67^, which in turn correlates with a regulation of mechanical forces^1^. By using traction forces microscopy and Shannon entropy correlation analysis, we were able to quantitatively show that CRB3 is needed to fine regulate the amount and alignment of forces. These data are in line with the fact that during collective motion epithelial cells increase and align their forces^20, 22^, however, similar behavior during the breaking of symmetry initiating epithelial transition was not described so far.

To our knowledge, this is the first study that clearly links a polarity protein, CRB3, to the remodeling and reorganization of the actin cytoskeleton and focal adhesions to the mechanical regulation leading to the breaking of symmetry during the shift between static to migrating epithelium, a key process in cell, developmental biology and mechanobiology.

## Supplementary Figures

**Fig. Supplementary 1:**
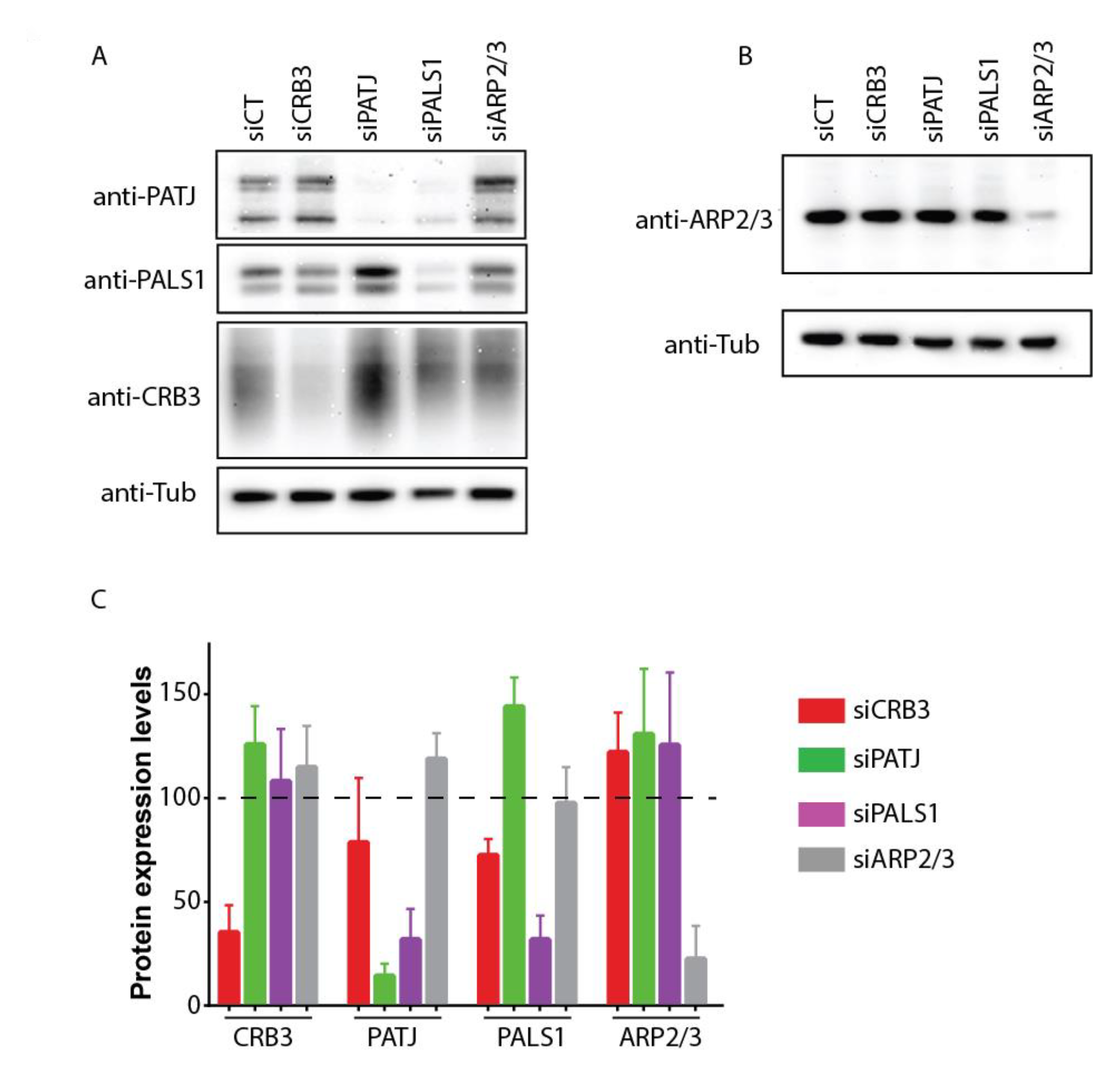
Protein expression levels after siRNA transfections. A-B) Immunoblot analysis showing the expression of PATJ, PALS1, CRB3 and Arp2/3, 5 days after transfections with siCT, siCRB3, siPATJ, siPALS1 and siARP2/3. α-Tubulin was used to standardize the loading conditions between the different depletions. C) Quantification of protein expression levels normalized to siCT cells. Data are represented as mean ± SEM. For each protein, n=3 samples pooled from 3 independent transfections.

**Fig. Supplementary 2:**
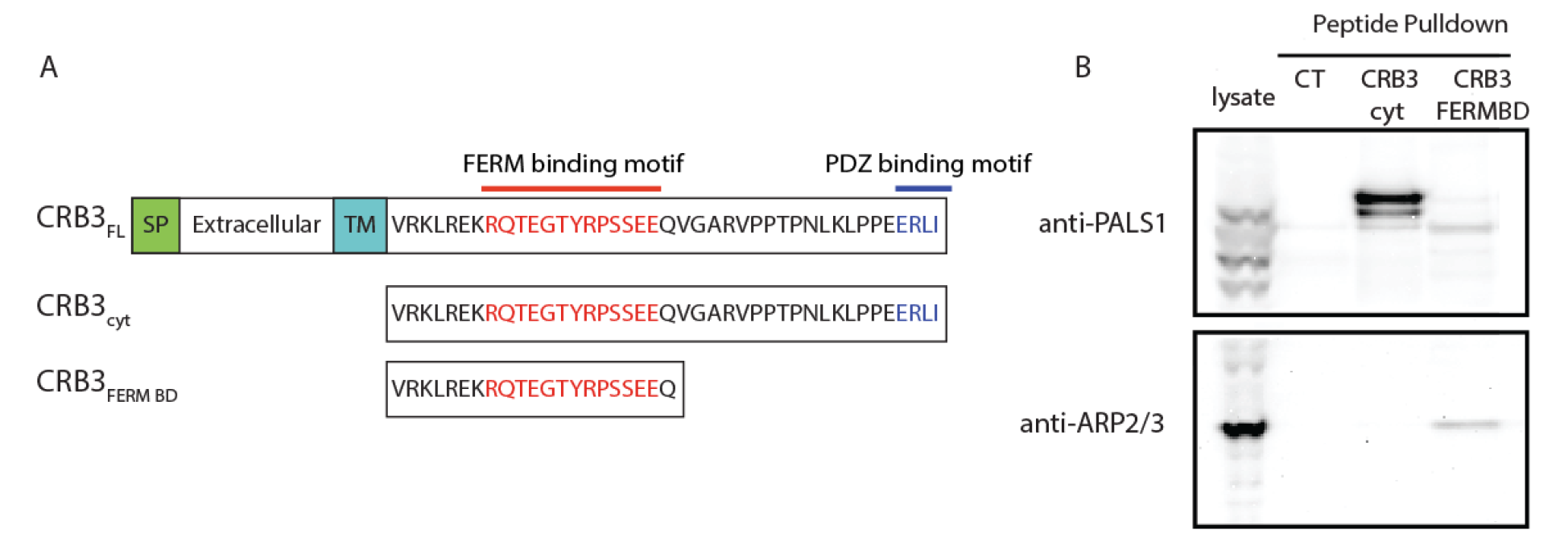
CRB3 interacts with ARP2/3. A) Scheme of CRB3 full length (CRB3 FL), cytoplasmic domain of CRB3 (CRB3 cyt) and the FERM binding domain of CRB3 that are used as peptide bait for the peptide pull-down. B) siCT lysates were pulled-down with the 3 peptides (CRB3 FL, CRB3 cyt and CRB3 FERMBD), and probed to detect PALS1 and ARP2/3 expression.

**Fig. Supplementary 3:**
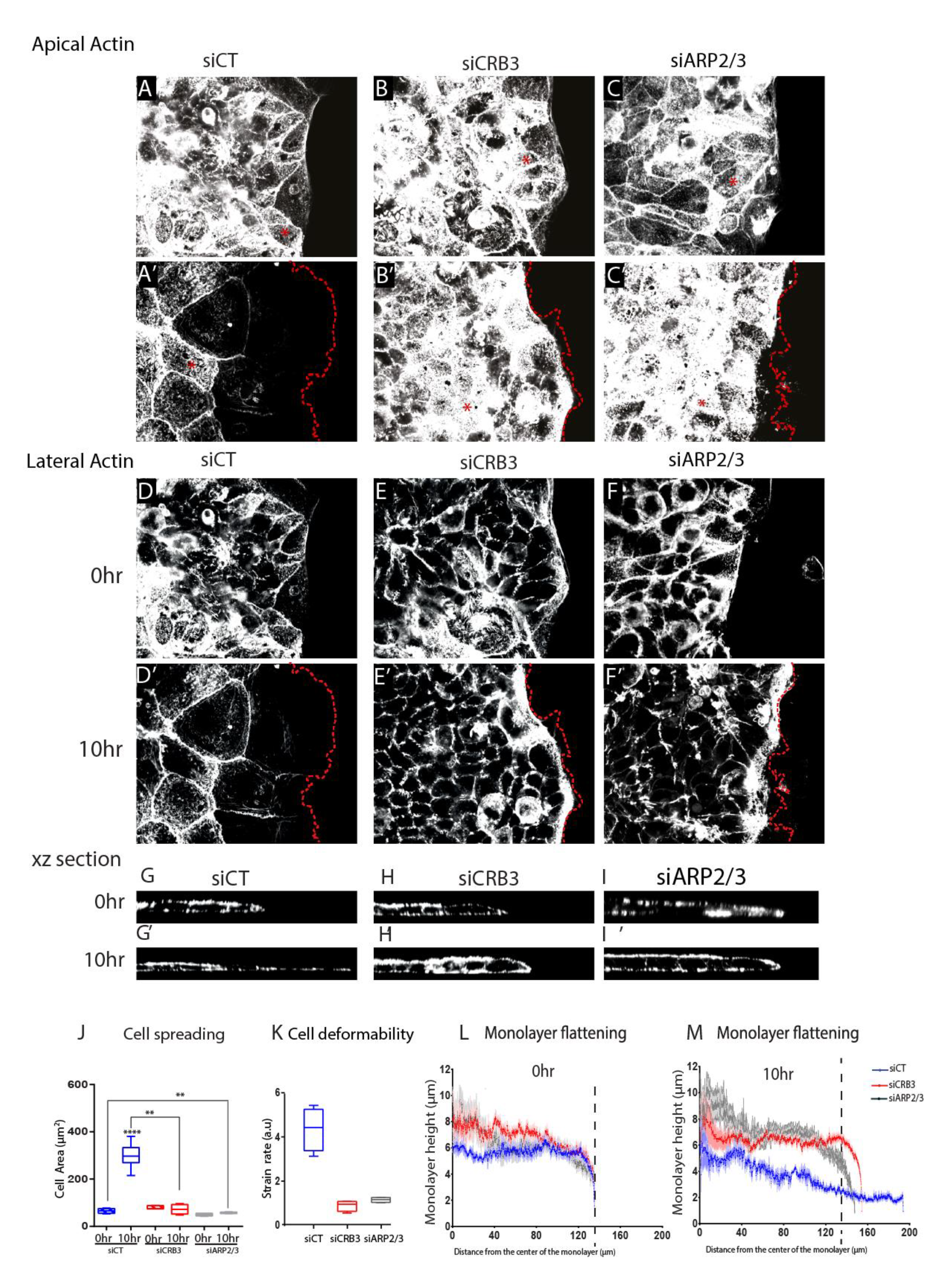
Downregulation of the CRB3 complex and ARP2/3 alter the organization of the F-actin. A-I’) Representative images showing the effect of each siRNA on the organization of the F-actin at the apical domain: siCT (A, A’), siCRB3 (B, B’), siARP2/3 (C, C’); at the lateral domain siCT (D, D’), siCRB3 (E, E’), siARP2/3 (F, F’), and in xz section siCT (G, G’), siCRB3 (H, H’) and siARP2/3 (I, I’) at 0hr and 10hr of migration. Red dash line: outline for the basal edge, red asterisk: microvilli. Scale bars= 15 µm. J-K) Quantification of cell spreading (J) and deformations (K) for each siRNAs at 0hr (dashed box plots) and 10hr (empty box plots) of migration. siCT, blue, n = 4 monolayers; siCRB3, red, n= 4 monolayers; siARP2/3, gray, n =4 monolayers. Data are presented as mean ± SEM. L-M) Monolayer height as function of the distance from the center of the cell monolayer at 0hr (L) and 10hr (M). For the different time points several lateral views were measured for each siRNAs (0hr, n=44, 10hr, n= 44, siCT, blue; siCRB3, red; siARP2/3, gray. Data are presented as mean ± SEM.

**Fig. Supplementary 4:**
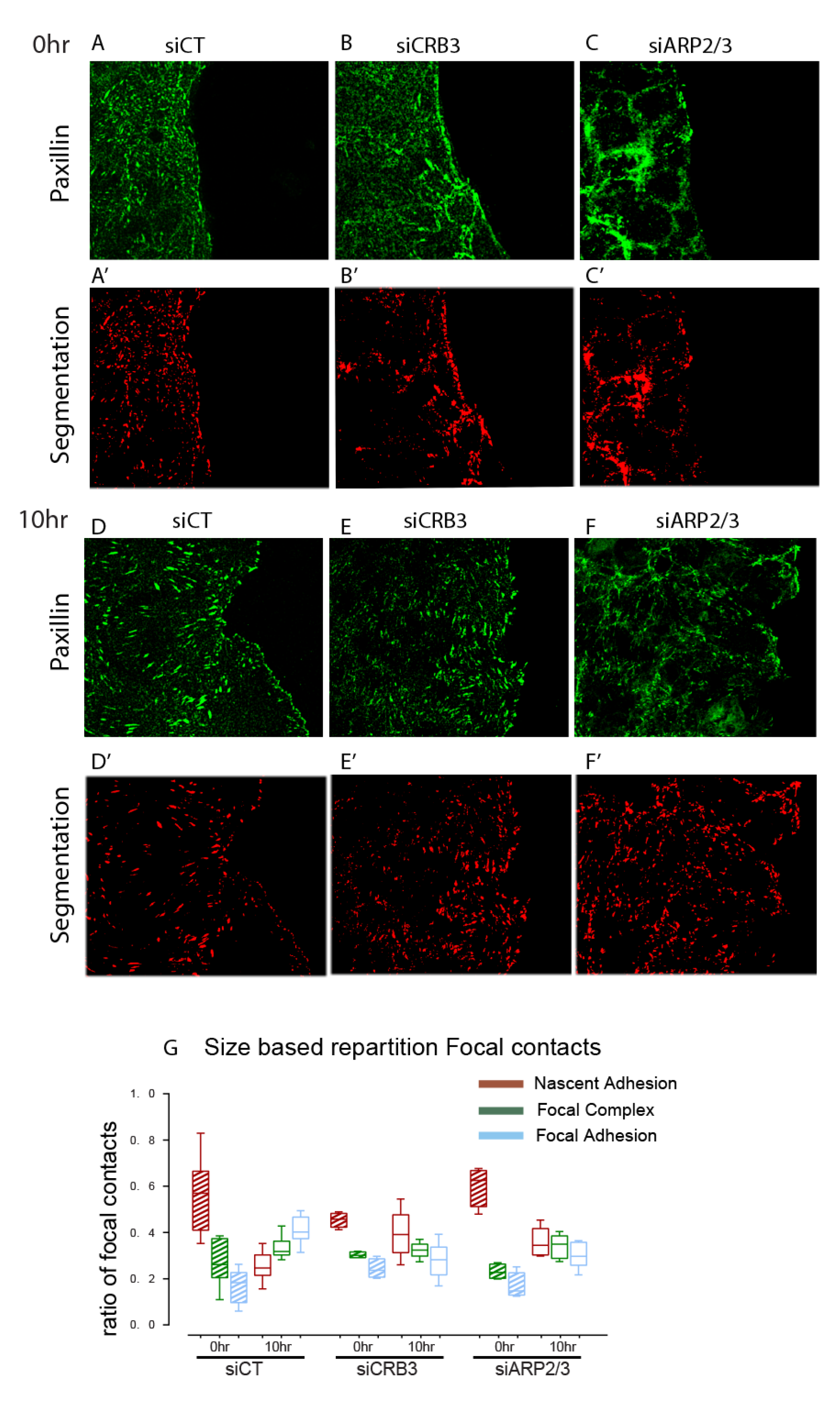
Downregulation of the CRB3 complex and ARP2/3 alter the organization of the focal adhesions. A-F’) Representative images showing the effect of each siRNA on the organization of the focal adhesion and the respective segmentation map: siCT (A, A’), siCRB3 (B, B’), siARP2/3 (C, C’) at 0hr and siCT (D, D’), siCRB3 (E, E’) and siARP2/3 (F, F’) at 10hr. G) Proportion of focal contacts as function of their sizes. The box plots represented the ratio between the number of focal contacts categorized by their sizes (0.25µm: brown, 0.5µm: green and >1µm: blue) to the total number of focal contacts at 0hr (dashed boxplot) and 10hr (empty box plot). The total number of focal contacts quantified for siCT, blue, n = 2530, 6 fields of view, 0hr, n= 6652, 15 fields of view, 10hr; siCRB3, red, n=2230, 4 fields of view, 0hr,n=7410, 17 fields, 10hr; siARP2/3, grey, n=1476, 4 fields of view, 0hr, n =5806, 13 fields of view, 10hr. Data are presented as mean ± Min Max.

## Material and Methods

### Caco2 cell line culture, transfection and drug treatment

Caco2 cells were grown on DMEM-glutamax media supplemented with 20% Foetal Bovine Serum, 1% of Non-Essential Amino Acid, 100U/mL Penicillin,Streptomycin.

siRNA transfections were performed by electroporation using Amaxa technology. 100pmoles of a pool of 3 siRNAs and 1.5 10^6^ freshly trypsinized Caco2 cells were transfected using the B24 program following the manufacturer recommendations. Cells were then seeded on a 6-well plates. 4 days after transfections, cells were trypsinized, 50000 cells were seeded on soft polyacrylamide gels and the remaining cells were processed by western blot.

The inhibition of branched actin was performed by adding 67µM, directly in the cell medium, of CK666 drug directly after the removal of the PDMS membrane.

### Preparation of soft polyacrylamide gels

Polyacrylamide gels with a Young’s modulus of 12 kPa were prepared as described previously (Bazellieres et al 2015). A solution of 19% acrylamide, 8% bis-acrylamide, 0.5% ammonium persulfate, 0.1% tetramethylethylenediamine, 0.64% of 200nm diameter red fluorescent carboxylate-modified beads and 2mg/mL NH-acrylate was prepared and allowed to polymerize at RT for 1hour. After polymerization, gels were incubated with 0.1mg/mL of collagen I overnight at 4°C.

### Fabrication of PDMS membrane

Polydimethylsiloxane (PDMS) membranes were fabricated according to procedures described previously (Bazellieres et al 2015). SU8-50 masters containing rectangles of 300×2,500 μm were obtained using conventional photolithography. Uncured PDMS was spin-coated on the masters to a thickness lower than the height of the SU8 rectangular feature and cured overnight at 60°C. A thick border of PDMS was left at the edges of the membranes for handling purposes. PDMS was then peeled off from the master and kept in ethanol at 4°C until use.

### Epithelial cell monolayer patterning

To pattern the cells on top of the polyacrylamide gels, a PDMS membrane was deposited on top of the polyacrylamide gel and 50000 cells were seeded within the rectangle defined by the PDMS stencil. Cells were allowed to adhere and differentiated on the gel for 24 hours. 15 minutes before time lapse imaging, the PDMS membrane was carefully removed allowing the cells to migrate toward the freely available substrate.

### Time lapse imaging and monolayer expansion quantification

Multidimensional acquisitions were performed on an automated inverted microscope (Zeiss AxioObserver, 10× lens) equipped with thermal, CO2, and humidity control, using Zen software. Images were obtained every 10 minutes during 600 minutes. 6 independent monolayers were imaged in parallel using a motorized XY stage.

Based on the phase contrast images, a segmented imaged of the edges of the monolayer was created at each time frame. The Euclidean distance between the two edges, at each pixel, for each siRNA was analyzed using a custom written MatLab codes based on the bwdist function.

### Immunoprecipitation and western blotting

For protein expression level confluent caco2 cells were lysed in lysis buffer (50 mM Tris HCL pH 7.4, 150 mM NaCl, 0.5% NP40) with protease inhibitor cocktail containing 1 μg/ml antipain, 1 μg/ml pepstatin, 15 μg/ml benzamidine, and 1 μg/ml leupeptin. Lysates were cleared by centrifugation at 20000g for 30min at 4°C. Protein expression levels were measured using Western Blot. Cell lysates were then mixed with Laemmli 1X and heated at 95°C for 5 minutes. Next, cell lysates were loaded to NuPAGE 4-12% Bis-Tris gel (ThermoFisher Scientific, Courtaboeuf, France) for electrophoresis. Proteins were then transferred to a nitrocellulose membrane (Whatman, GE Healthcare Life Sciences), which was blocked with 5% dry-milk-Tris Buffer saline, 0.2% Tween, and incubated with primary antibodies (overnight at 4°C) followed by the horseradish peroxidase coupled secondary antibodies (1h, room temperature). Bands were revealed using chemi-luminescence reagent plus (Perkin Elmer) and visualized by MyECL imager (ThermoFisher Scientific, Courtaboeuf, France). The intensity of the bands was quantified using ImageJ software. Tubulin was used as an endogenous control for normalization. Protein concentrations are reported relative to the control.

A phase separation method was used to quantify the amount of ARP2/3 that is associated at the cell membrane in siCT and siCRB3 conditions. Confluent caco2 cells were lysed in lysis buffer (50 mM Tris HCL pH 7.4, 150 mM NaCl, 2% Triton X-114) with protease inhibitor cocktail containing 1 μg/ml antipain, 1 μg/ml pepstatin, 15 μg/ml benzamidine, and 1 μg/ml leupeptin. Lysates were cleared by centrifugation at 20000g for 30min at 4°C, and the supernatant was brought up to 37°C for 5min allowing to collect the phase that contains the membrane bound proteins ^68^. After 4 washes with lysis buffer without Triton-X114, bound proteins were processed by western blotting.

CRB3 peptide pull-down assays: 150 µL pelleted streptavidin beads (streptavidin agarose resin, ThermoFisher Scientific, Courtaboeuf, France) were coated with 2mg of a biotinylated peptide mimicking the cytoplasmic part of CRB3A (amino-acids 90 to 120) or with the cytoplasmic part containing the FERM binding domain (amino-acids 90 to 100) called CRB3 FERMBD, all synthetized by CovaLab (Cambridge, UK). The cell lysates were centrifuged at 20 000g during 30 min and the supernatants were incubated with 20 µl of streptavidin beads CRB3cyt or CRB3 FERMBD at 4°C overnight. After 4 washes with lysis buffer, bound proteins were processed by western blotting.

### Quantification of activated Rac and Rho

Caco2 cells siCT and siCRB3 were lysed in lysis buffer (50mM Tris pH 7.4, 500mM NaCl, 10mM MgCl_2_, 1% Triton X-100, 0.1% SDS, 0.5% Sodium Deoxycolate) with protease inhibitor cocktail containing 1 μg/ml antipain, 1 μg/ml pepstatin, 15 μg/ml benzamidine, and 1 μg/ml leupeptin, for 5min on ice. Cell lysates were centrifugated and the supernatants were incubated with the GST fused to Rhotekin-p21Binding Domain (GST-Rhot-PBD) or PAK-p21Binding Domain (GST-PAK-PBD) at 4°C for 30 minutes. The constructs were kindly provided by Michael Sebbagh, CRCM, Marseille. After 3 washes with lysis buffer, bounds proteins were processed by western blotting.

### Immunofluorescence

Caco2 cells were washed with PBS, fixed with 3% paraformaldehyde for 10 minutes and permeabilized in 0.5% triton X-100 for 5 minutes. Cells were blocked in 10% FBS for 1 hour at room temperature before being incubated for 4 hours, room temperature, with primary antibodies. After incubation with the appropriate fluorescence-conjugated secondary antibodies for 1hr at room temperature, cells were washed and mounted in DABCO/Mowiol mounting media. Images were acquired with a Zeiss 510 meta confocal microscope, using a 63× Objective with 1.4 NA lens.

### Quantification of actin properties

Images were taken using a Zeiss 510 Meta confocal microscope (Zeiss) and were analyzed as following. Actin orientation and coherency were measured using the OrientationJ plugin, the vector field to compute the different parameters was extracted using a grid size of 30 pixels and an α value of 2 pixels. The orientation was defined as the orientation index such as OI =cos (Ɵ), Ɵ defining the angle between the axis of the actin fiber and the direction of migration.

### Quantification of cell and focal adhesion properties

Images were taken using a Zeiss 510 Meta confocal microscope (Zeiss) and were analyzed as following. Cell contours were detected using cortical actin fluorescent signals, monolayer heights were detected using Z confocal section using actin staining and focal contact were detected using paxillin fluorescent signals in the immunostaining images. Using custom written Matlab codes based on the regionprops function, all the cell contours, monolayer heights and focal contacts within an image were automatically segmented and localized.

Based on the segmented images obtained for the cell contours and focal contacts, an ellipse was fitted on each feature and different parameters were extracted such as ellipse area, length of the major and minor axis. The ellipse orientation was defined as the orientation index such as OI =cos (Ɵ), Ɵ defining the angle between the major axis and the direction of migration.

Based on the segmented images obtained for the monolayer height, the Euclidean distance between the two edges, at each pixel, for each siRNA was analyzed using a custom written MatLab codes based on the bwdist function.

### Quantification of focal adhesion dispersion

The centroid of each focal contact was automatically determined from the segmented images obtained previously with custom written Matlab scripts. XY position of all centroids were used to build triangles between the nearest neighbors with the Delaunay Triangulation Matlab script. Once the triangulation was obtained, the areas of the triangles were calculated in Matlab.

### Traction Force Microscopy

Traction forces were computed using Fourier Transform Traction Microscopy with finite gel thickness. Briefly, as cells migrate, they exert force on the underlying the substrate. Gel deformations are observed by imaging the fluorescent beads embedded within the gels. Gel displacements between any experimental time point and a reference image obtained after cell trypsinization were computed using particle imaging velocimetry software. To reduce systematic biases in subpixel resolution and peak-locking effects, we implemented an iterative process (up to four iterations) based on a continuous window shift technique (Serra-Picamal et al., 2012). Traction vectors Ti,j(t) within the field of interest are obtained from displacement vectors ui,j (t) for all time points t = 1,…, n and locations (i,j) of the M × N gel interface matrix.

### Monolayer Stress Microscopy

Maps of inter-and intracellular tension within the monolayer were computed using monolayer stress microscopy. In a 2D approximation, monolayer stress is fully captured by a tensor possessing two independent normal components (σxx and σyy) and two identical shear components (σxy and σyx). At every pixel of the monolayer, these four components of the stress tensor define two particular directions of the plane, one in which the normal stress is maximum and one in which it is minimum. These directions, which are mutually orthogonal, are called principal stress orientations, and the stress values in each principal orientation are called maximum (σ11) and minimum (σ22) stress components. The average normal stress is defined as σn = (σ11+σ22)/2. The spatial resolution and force precision of MSM are formally set by those in the original traction maps.

### Shannon Entropy Analysis

To apply Shannon’s entropy, we partition the range of the angle represented as the angle between two dimensional vectors, into 30° angle bins histogram. With this histogram, the probability of angles in each bin can be calculated, and the information content can be computed using Shannon’s entropy as

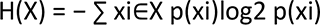

The Shannon’s entropy allows to measure and thus compare the amount of variation in angle between the vectors in siCT, siCrb3 and siARP2/3 conditions. When the angles between the vectors are different (less aligned) the number of information is high (1), whereas an alignment of the vectors tends to have a more deterministic number of information (0).

### Computation of z-scores

The z-score is defined as the signed number of standard deviations an observed quantity deviates from the mean of that quantity. In our study the the z-score of a quantity x (a physical or biological property) in response to a siRNA perturbation is defined as:

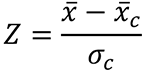

Where 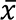 is the mean of x under the siRNA perturbation, 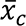 is the mean of x under control conditions, and 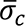 is the standard deviation of x under control conditions.

### siRNAs, Antibodies and reagents

The siRNA sequences used were: siCRB3 (5’-GCAAAUACAGACCACUUCU-3’, 5’- CUGCUAUCAUCGUGGUCUU-3’, 5’-GUGCGGAAGCUUCGGGAGA-3’, 5’-GCUUAAUAGCAGGGAAGAA-3’, Dharmacon (On-Target plus Smart Pool)), siCT (5’- CGUACGCGGAAUACUUCGAtt-3’, Ambion), siPALS1 (5’-UUCCUUAUGAUGAACUGGCtt-3’) and siPATJ (5’-CCAGAUACUCACACUUCAGtt-3’, Ambion), siARP2/3 (5’- GGAUUCCAUUGUGCAUCAAtt-3’, 5’-GGGAUGAUGAGACCAUGUAtt-3’, 5’- AAAUCCUAAUGGAGACAAAtt-3’, Ambion)

The following primary antibodies were used : rabbit anti CRB3 D2 (Lemmers et al 2004), rat anti CRB3 1E6 (MABT1366 Merck), mouse anti-paxillin (BD transduction 612405), rabbit anti-PATJ Ina2 (Lemmers et al, 2002), chicken anti-PALS1 SN47II (gift from Jan Wijnholds, Kantardzhieva et el, 2005), mouse anti-PALS1 (MPP5, Abnova H00064398), rabbit anti-p34- ARPC2 (07227I, Sigma Aldrich), rabbit anti-pEzrin (Abcam ab47293), mouse anti-Rac (BD transduction, Clone 102), mouse anti-RhoA (SantaCruz 26C2, sc 418). The secondary antibodies used were : Alexa Fluor 488 anti-rat (Invitrogen, A21208), Alexa Fluor 488 anti mouse (Invitrogen, A21202), Alexa Fluor 647 mouse (Jackson Immuno Research 715-605-151) Alexa Fluor 488 anti-rabbit (Invitrogen, A21206), and HRP anti-rat (Jackson Immuno Research 712 035 153), HRP anti-mouse (Jackson Immuno Research, 715 035 151), HRP anti-rabbit

(Jackson Immuno Research 111 035 003), HRP anti-chicken (Jackson Immuno Research 703 035 155). The following probe was used for actin labeling: Phalloidin 647 (Cell Signalling technology, 8940S).

## Abbreviation

ABP: (apico-basal polarity)
EMP: (Epithelial to Mesenchymal Plasticity)
CRB3: (Crumbs3)
PALS1: (Protein Associated with Lin Seven 1)
PATJ: (PALS1-Associated Tight Junction)
ARP2/3: (Actin Related Protein 2/3 complex)
KD: (knockdown)
PDMS: (Polydimethylsiloxane)

## Acknowledgments

We would like to thank Andrea Pasini, Maria Mandela Prunster and Delphine Delacour for discussion.

## Funding

DMH salary is from INSERM, ALB and EB salaries are from CNRS, NG salary is from the Turing Center for Living systems (CenTuri) (ANR-16- CONV-0001).

We acknowledge the IBDM imaging facility, member of the national infrastructure France-BioImaging supported by the French National Research Agency (ANR-10-INBS-04). This work was supported by CNRS and AMU (UMR7288), by the CapoStromex project from the A*MIDEX project (ANR-11-IDEX-0001-02) and the LabEx INFORM (ANR-11-LABX-0054) funded by the “Investissements d’Avenir” French Government program.

## Author contributions

EB conceived the study and designed experiments. EB, DMH, NG performed the experiments, EB and DMH analyzed data. EB developed data analysis tools. VC developed computational mechanical tools. EB and ALB wrote the manuscript. All authors discussed the results and contributed to the manuscript.

